# Cytotoxic T-cell-based vaccine against SARS-CoV2: a hybrid immunoinformatic approach

**DOI:** 10.1101/2021.09.26.461851

**Authors:** Alexandru Tîrziu, Virgil Păunescu

## Abstract

This paper presents an alternative vaccination platform that provides long-term cellular immune protection mediated by cytotoxic T-cells. The immune response via cellular immunity creates superior resistance to viral mutations, which are currently the greatest threat to the global vaccination campaign. Furthermore, we also propose a safer, more facile and physiologically appropriate immunization method using either intra-nasal or oral administration. The underlying technology is an adaptation of synthetic long peptides (SLPs) previously used in cancer immunotherapy. SLPs comprising HLA class I and class II epitopes are used to stimulate antigen cross-presentation and canonical class II presentation by dendritic cells. The result is a cytotoxic T cell-mediated prompt and specific immune response against the virus-infected epithelia and a rapid and robust virus clearance. Peptides isolated from COVID-19 convalescent patients were screened for the best HLA population coverage and were tested for toxicity and allergenicity, 3D peptide folding, followed by molecular docking studies provided positive results, suggesting a favourable antigen presentation.

## Introduction

SARS-CoV2 is an RNA virus responsible for the pandemic that we are facing in present. COVID-19 clinical features depend on the genetic variants of both the patient and the virus, inoculum size, and the presence of comorbidities. [1] The main transmission routs are respiratory and oral [2], suggesting the importance of the mucosal immunity in disease onset. No specific antiviral medication is available [3], so prevention is the key for reducing the mortality associated with COVID-19.

Results from previous studies related to previous endemic outbreaks involving SARS-CoV and MERS-CoV suggested that coronaviruses trigger T cell and antibody immune responses in infected patients, but the antibody levels seem to become undetectable 2-3 years after recovery [4], while SARS-CoV-specific memory T-cells were identified after more than a decade after the infection. Ng et al. isolated in 2016 memory T cells 11 years after the infection, while le Bert et al. in 2020 discovered a more than 17 years viral-specific cellular immune response. [1]

Analysis of the cellular and immune response from COVID-19 convalescent patients revealed that those with mild COVID-19 presented a vigorous T-cell mediated response months after SARS-CoV2 infection and a low to undetectable titre of antibodies. [5] In contrast, severe COVID-19 presented an early phase lymphocytopenia followed by intense cytokine release and high antibody titres, due to poor viral clearance. [6] In the study performed by Sekine et al, 28% of healthy individuals presented cross-reactive memory T-cells against SARS-CoV2. These findings confirm the bipolar role of T cells in COVID-19 pathogenesis – a higher number of lymphocytes during the early phase of infection assures a rapid and efficient antiviral response, whereas an early lymphocytopenia, followed by a subsequent immune hyperactivation leads to a poorer prognosis.

Lymphocytopenia is one of the most decisive factors in the evolution of COVID-19. Resolved lymphocytopenia predicts a favourable outcome, while unresolved lymphocytopenia leads to a poor prognosis. T lymphocytes were shown to hamper the innate immune response and prevent immune hyperactivation. [10] Thus, lymphocyte priming is a desirable event in the context of vaccination.

Mucosal immunity stimulation represents a promising alternative to traditional vaccination due to its simplicity, acceptance rate (less unpleasant than injections) and a reduced number of post-vaccinal complications. By targeting the pathogen right at the entry site, the viral replication decreases substantially and the risk of evolution to more severe disease forms is mitigated.

In present, vaccination platforms range from live attenuated formulas to the newly introduced mRNA and viral vector-based platforms. Peptide-based vaccination provide a promising alternative due to its high specificity, biological activity and tissue penetration as well as low production costs. [11,12]

Compared to single epitopes or protein subunits, SLPs present several advantages such as reduced CTL (cytotoxic T lymphocyte) tolerance, higher stability, T helper cell involvement and enhanced peptide repertoire.

Single class I-restricted epitopes begin to diffuse systemically upon administration and bind randomly to HLA class I molecules expressed on all nucleated cells. Because non-antigen-presenting cells lack costimulatory proteins, CTLs cannot be activated and immune tolerance is induced. [12,15]

In contrast, by expanding the CD8+ epitope with a class II-restricted epitope, the resulting amino acid chain will be forced to be internalized and processed by the antigen-presenting cells. A proportion of peptides will be processed in the endosomal pathway and expressed with the HLA class II molecule, while others enter the vacuolar pathway to be cross-presented with the HLA class I molecule. [16]

In terms of stability, single CD8+ epitopes bound to the MHC class I molecule are expressed for a short amount of time, leading to a weak and transient immune response. On the other hand, SLP-derived class I epitopes expressed higher stability when cross-presented.

By including a class II-restricted epitope, T helper cell stimulation is produced with subsequent cytokine release and immune response augmentation. [12]

SLPs also have the advantage of combining peptide sequences located at distant sites inside the same protein or amino acid chains originating from different proteins. As a result, the immunogenic peptide repertoire is increased, providing a more specific immune response. [15]

By expanding the peptide repertoire, SLPs can also function as immune response enhancers. Coppola et al. showed that administration of synthetic long peptides derived from Mycobacterium tuberculosis Latency Antigen Rv1733c, a protein expressed in dormant bacteria, led to an improved bacterial clearance in HLA-DR3 transgenic mice. [17]

Due to their high data processing and analysis capacities, in silico methods can be a useful tool for characterizing diverse immunological events and speeding up the process of vaccine design. Immunoinformatic and computational biology approaches were already used for designing vaccines against infectious agents such as Helicobacter pylori[18], Vibrio cholerae [19], Plasmodium species [20] or yellow fever virus. [21] For COVID-19, research groups designed multi-epitope subunit protein vaccines [22,23], but this vaccine landscape lacks a SLP-based vaccine design.

Our group previously published an in-silico approach to generate SLPs against SARS-CoV-2 infection for the Romanian population, based on regional phenotypic characteristics. [19] Following the previous study, the current technology uses a hybrid approach based on immunoinformatic methods described in the previous article, as well as data collected from databases containing epitopes from COVID-19 convalescent patients.

## Methods

### Epitope screening

Epitopes were screened from a peptide pool comprising 1209 peptide sequences identified in 852 patients recovered from COVID-19. [25] The peptide pool database can be accessed here: https://www.mckayspcb.com/SARS2TcellEpitopes/. Main screening criteria included a high response frequency, a high degree of conservation (>0.85) and cross-specificity (to cover multiple HLA alleles).

### Population coverage analysis

The peptide sets were screened using IEDB population coverage tool, an algorithm that calculates the percentage of a population of interest that will be covered by a user defined peptide-HLA dataset. [26] Further selection was performed so that the peptide pool would cover the maximum percentage of population with the minimum number of peptides.

### Synthetic long peptide construction

As previously described by Rabu et al., the synthetic long peptide construct with a higher probability to be presented and cross-presented to T cells by the dendritic cells comprises a HLA class II-restricted epitope at the N-terminus, a 6-mer cathepsin-sensitive linker sequence (LLSVGG) and a HLA class I-restricted epitope. [15]

### Allergenicity screening

Allergenicity testing was performed using AllerCatPro, a webserver which compares the query structure with FASTA sequences and 3D structures from an extensive database of allergens. In this manner, both linear and discontinuous allergenic epitopes are detected. [27]

### Toxicity screening

Toxicity was assessed using ToxinPred webserver (http://crdd.osdd.net/raghava/toxinpred/), a support vector machine algorithm that separates non-toxic peptides from the toxic peptides based on a training dataset from SwissProt, TrEMBL and other protein databases. [11]

### Three-dimensional structure prediction

Rosetta ab initio was used to predict the three-dimensional conformations for each synthetic long peptide. For a query peptide/ protein, Rosetta ab initio approximates its secondary and tertiary structure by using libraries of 3-mer and 9-mer secondary structures. Generation of 3-mer and 9-mer libraries is performed using Robetta by searching the most probable 3-mer and 9-mer conformations in already solved structures.

### Three-dimensional structure validation

3D structure validation was performed to assess whether the 3D structure prediction provided stable, good quality models. For this step we used PROCHECK[28] and their corresponding Ramachandran plot were drawn. 3D structure visualization was performed using PyMol.

### Molecular docking studies

To investigate how likely the SLPs are to be internalized by the antigen-presenting cells, molecular docking studies were performed using HADDOCK 2.4. [29,30] The antiviral innate immune receptors TLR2 and TLR4 (Toll-like receptor) were used for this assay. By binding to Toll-like receptors 2 and 4, the subsequent cytokine release can trigger the internalization of the SLPs. Toll-like receptor three dimensional structures were downloaded from the PDB database with the PDB ids 6NIG [31] and 3FXI [32] respectively.

The initial step for HADDOCK 2.4 molecular docking is it0 in which both the ligand and the receptor are treated as rigid objects in the tridimensional space. The algorithm searches for the most geometrically favourable surface for the ligand to bind to the receptor. The second step (it1) is a flexible docking in which the torsion angles from the active residues of both the ligand and the receptor are modified to produce strong physical intermolecular bonds (hydrogen bonds, ionic interactions etc.). The last step (itw) exploits the capacity of the ligand to displace the water molecules surrounding the active site once bound to the receptor. This refining process adjusts the torsion angles so that the SASA (solvent accessible surface area) is minimized. Further structure refinement was performed using HADDOCK 2.4 to reduce the RMSD, restraints violation energy and HADDOCK score minimization. Gibbs free energies and dissociation constants were calculated using PRODIGY web server. [33,34]

## Results

Nineteen peptides were identified based on the aforementioned criteria: 15 HLA class I-restricted and 4 class II-restricted. The most epitopes originated from the S protein (13/19, 68.42%) and M protein (4/19, 21.05%) probably due to their position on the viral surface membrane which facilitates antigen recognition. (Table 1) Most peptides are conserved throughout SARS-CoV2 variants, indicating that the existing SARS-CoV2 mutations do not influence the antigen presentation and immune response after peptide vaccination.

**Table 1.**
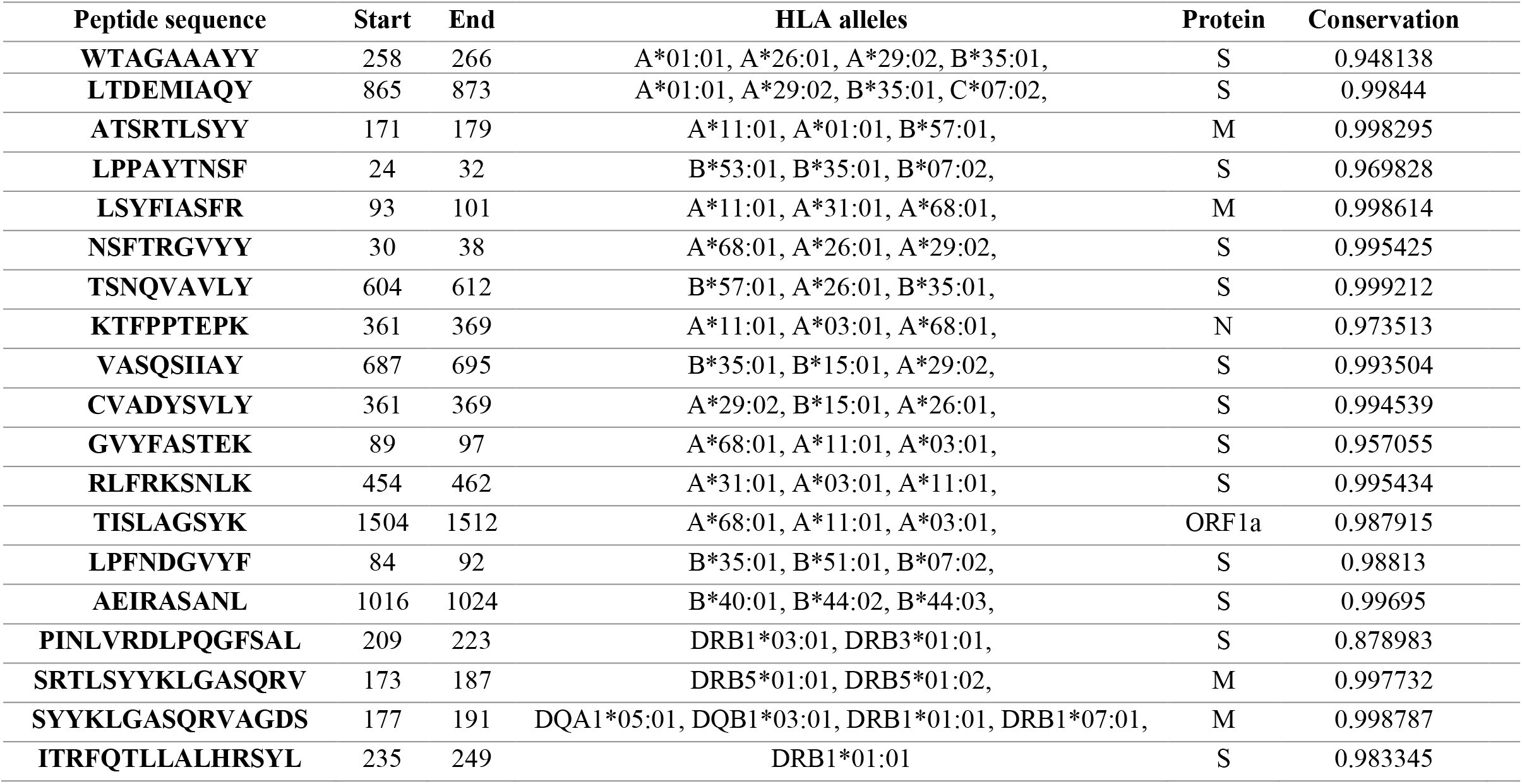
Selected HLA class I and class II-restricted epitopes based on their conservation and the number of allele hits.

Population coverage analysis showed that the class I coverage was 85.94%, class II coverage 75.42% and combined coverage 96.54%. The average number of epitope hits / HLA combinations recognized by the population was 4.49 for class I, 1.27 for class II and 5.76 for the combined set. PC90 for the combined set was 1.81, which roughly translates to a minimum number of 2 peptides that will be recognized by 90% of the population. (Figure 2)

**Figure 1.**
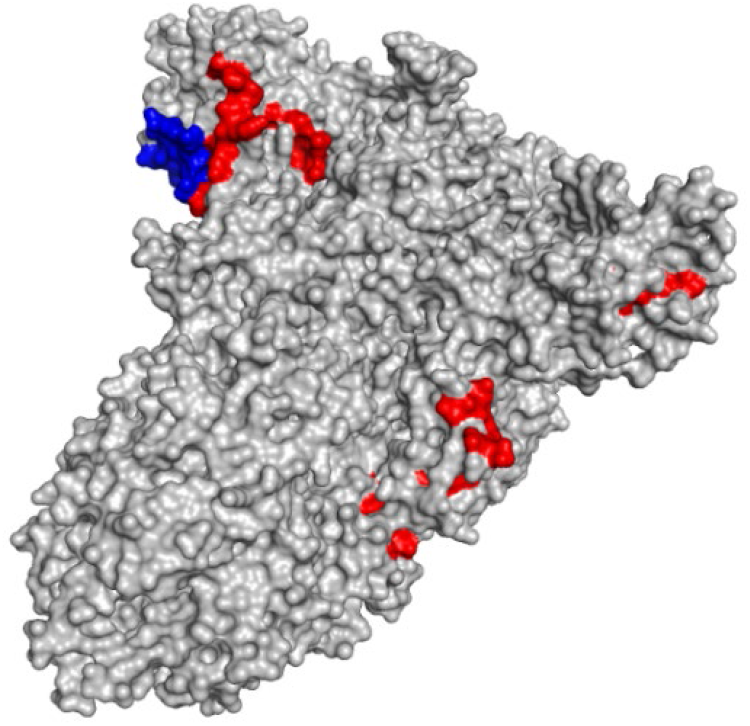
3D representation of the spike (S) protein (gray) with highlighted class I-restricted (red) and class II-restricted epitopes.

**Figure 2.**
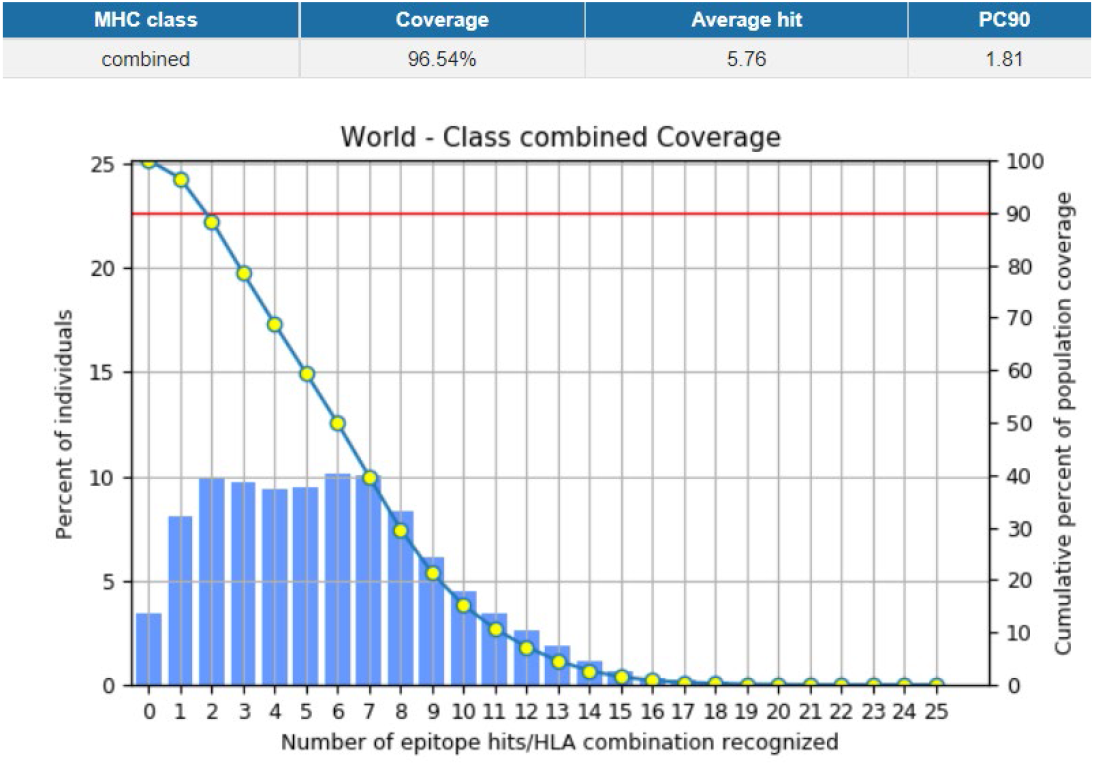
Population coverage analysis for the combined peptide set.

SLP construct comprises a HLA class II and a HLA class I-restricted epitope joined by a cathepsin-sensitive linker (LLSVGG). The choice of this linker was made based on the experimental data of Rabu et al. on in vitro and in vivo antigen presentation assays.

None of the peptides proved to be allergenic or toxic. (Table 2)

**Table 2.**
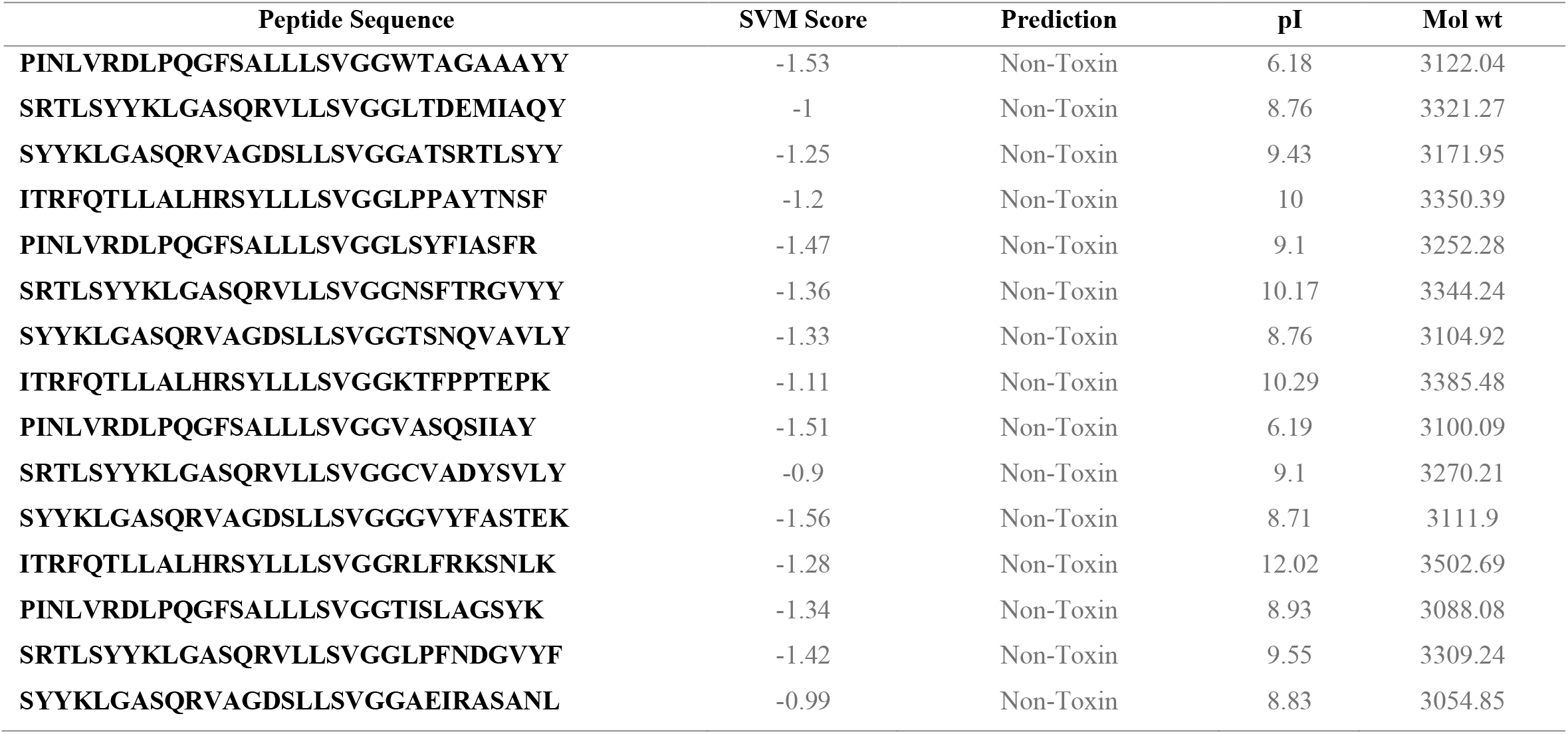
Predicted synthetic long peptide (SLPs) sequences, toxicity prediction, isoelectric point (pI) and molecular weight (Mol wt).

For each synthetic long peptide structure we predicted 100 three-dimensional models and ranked them based on the Rosetta score. The model with the best score was selected for further analysis. (Table 3) Ramachandran plots revealed that the percentage of residues in most favourable regions is below 90%, suggesting good quality model predictions. (Figure 3) The structures were then visualized with PyMol. (Figure 3)

**Table 3.**
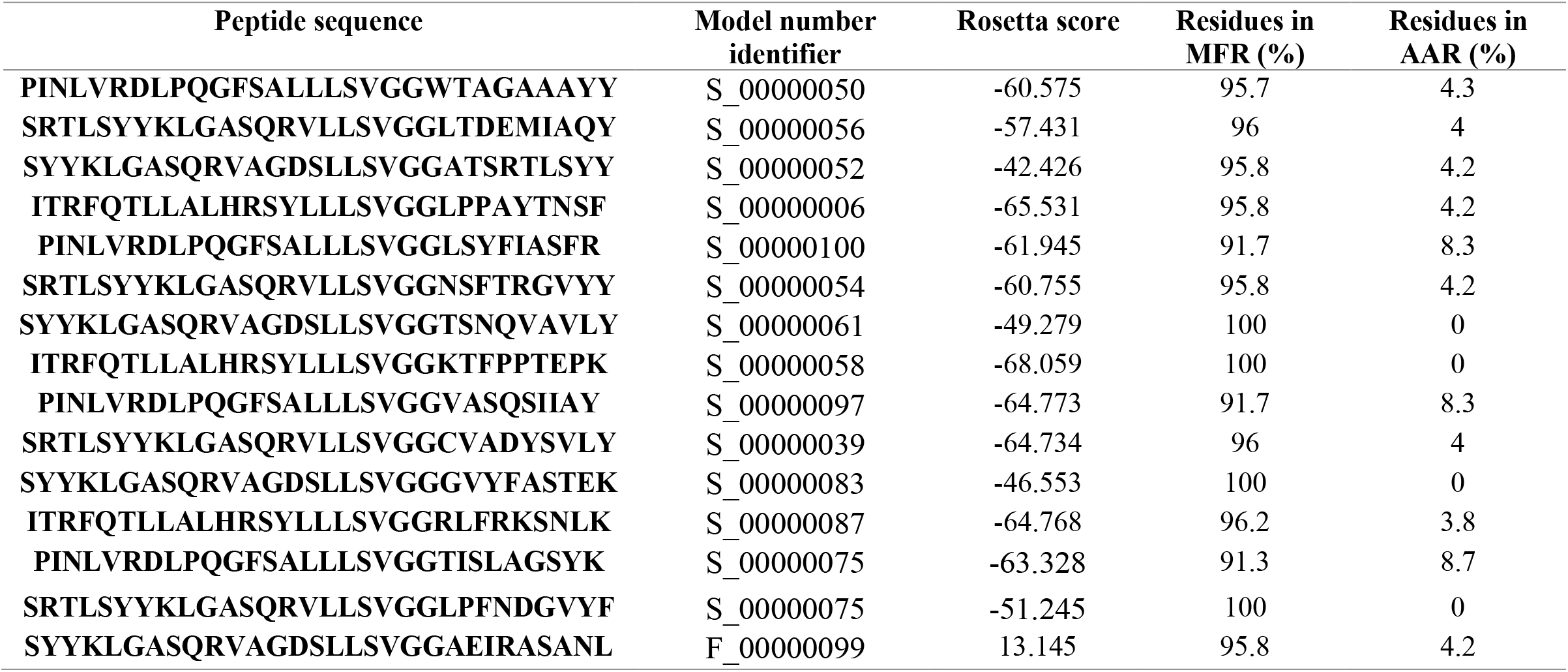
Model number for each synthetic long peptide and its Rosetta score. MFR – most favourable regions; AAR – additional allowed regions.

**Figure 3.**
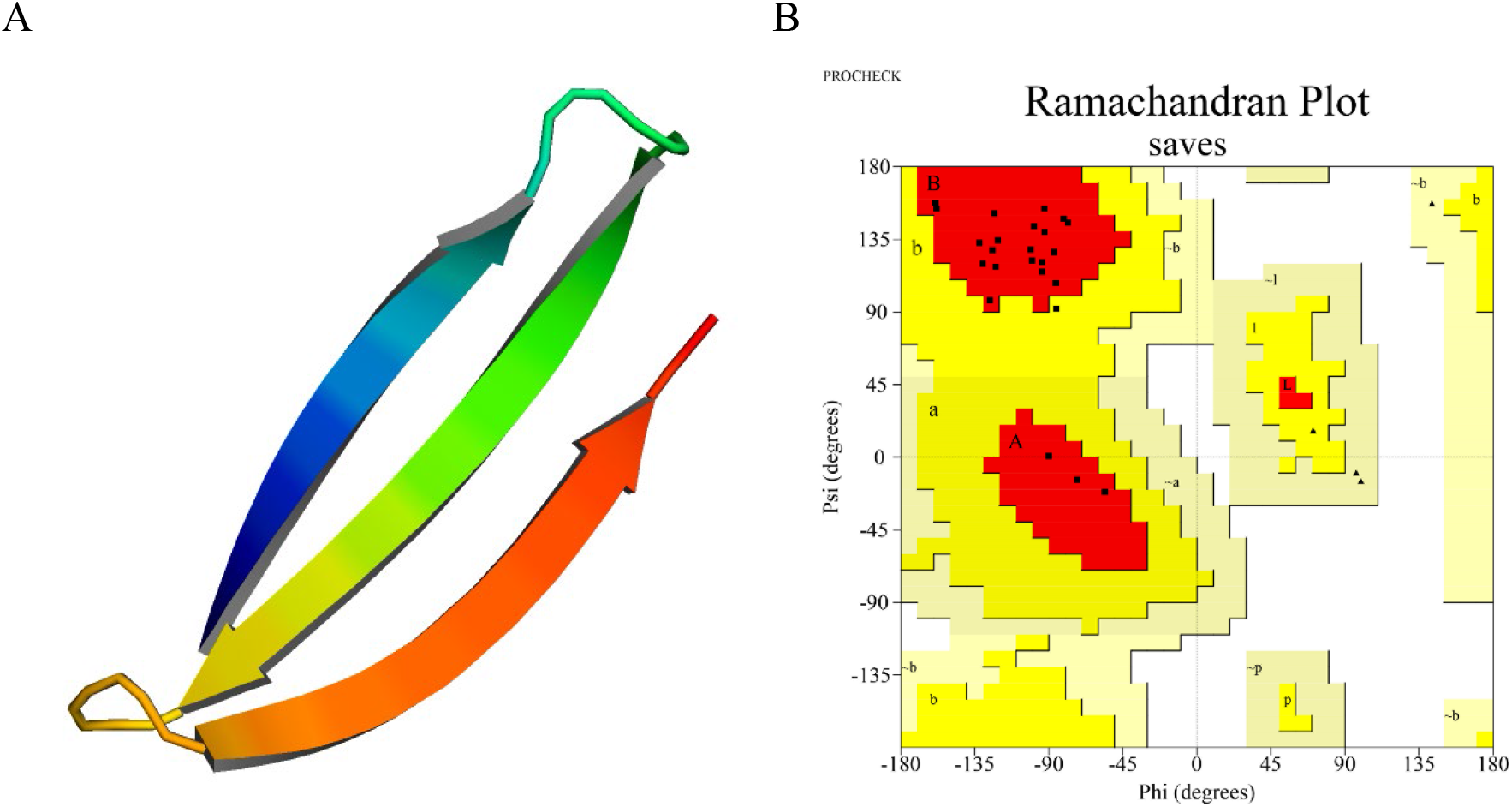
A. 3D structure visualization of the synthetic long peptide PINLVRDLPQGFSALLLSVGGWTAGAAAYY using PyMol. B. Ramachandran plot for the corresponding SLP.

**Figure 4.**
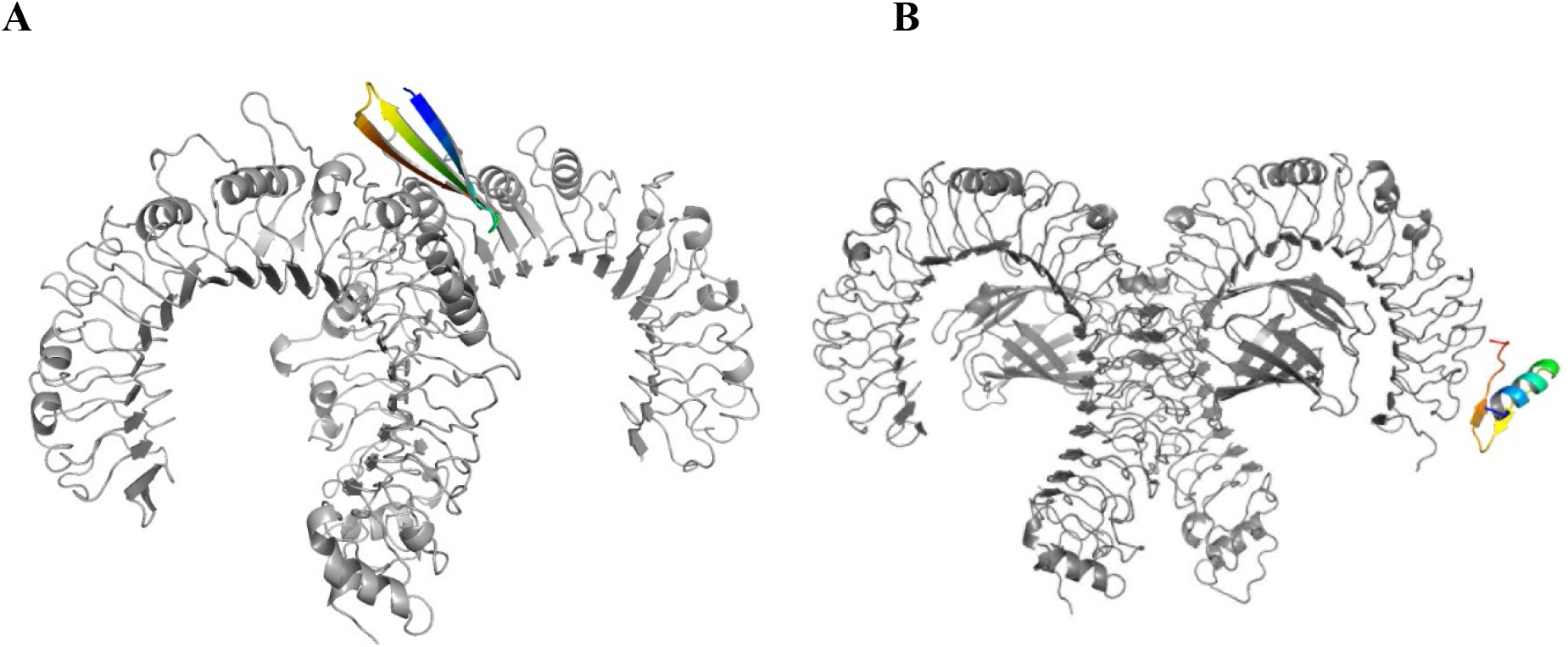
A. SLP (colored) bound to TLR2 (gray). B.Toll-like receptor 4 (gray) in complex with a synthetic long peptide (colored).

Molecular docking of the SLPs with TLRs was performed using HADDOCK 2.4. For each SLP, the predicted structures of the docked complex were grouped into clusters. The cluster with the best RMSD, van der Waals, electrostatic and desolvation energy values was selected for further analysis.

TLR-SLP complexes three-dimensional structures were refined using HADDOCK refinement tool. (Table 4) 3D structure visualization was performed using PyMol. (Figure 3) Gibbs free energy and dissociation constants indicate a highly probable interaction between the TLR2/4 and the designed synthetic long peptides. (Tables 4 and 5)

**Table 4.** HADDOCK results for TLR2-SLP molecular docking (after refinement).

| Peptide # | HADDOCK score | RMSD | Van der Waals energy | Electrostatic energy | Desolvation energy | Restraints violation energy | Buried Surface Area |
| --- | --- | --- | --- | --- | --- | --- | --- |
| 1 | -129.3 +/- 0.8 | 0.5 +/- 0.3 | -71.3 +/- 0.8 | -199.9 +/- 4.7 | -18.2 +/- 0.8 | 0.8 +/- 0.5 | 1824.8 +/- 33.9 |
| 2 | -116.6 +/- 2.8 | 0.5 +/- 0.3 | -60.5 +/- 2.1 | -316.9 +/- 22.8 | 7.2 +/- 1.3 | 0.6 +/- 0.1 | 1746.5 +/- 17.9 |
| 3 | -120.9 +/- 0.9 | 0.5 +/- 0.3 | -71.6 +/- 2.8 | -259.1 +/- 26.0 | 2.4 +/- 2.7 | 1.0 +/- 0.3 | 1912.0 +/- 16.8 |
| 4 | -105.7 +/- 1.8 | 0.5 +/- 0.3 | -65.3 +/- 1.2 | -155.5 +/- 9.2 | -9.3 +/- 3.3 | 0.5 +/- 0.1 | 1730.8 +/- 40.7 |
| 5 | -86.9 +/- 2.0 | 0.5 +/- 0.3 | -46.8 +/- 3.1 | -213.4 +/- 1.0 | 2.5 +/- 1.2 | 0.7 +/- 0.3 | 1352.3 +/- 43.7 |
| 6 | -100.2 +/- 2.2 | 0.5 +/- 0.3 | -56.9 +/- 2.6 | -221.2 +/- 9.9 | 0.9 +/- 1.2 | 0.4 +/- 0.1 | 1666.2 +/- 31.5 |
| 7 | -111.1 +/- 1.4 | 0.5 +/- 0.3 | -67.8 +/- 3.4 | -179.0 +/- 25.4 | -7.5 +/- 1.9 | 0.7 +/- 0.4 | 1830.4 +/- 44.8 |
| 8 | -123.4 +/- 3.7 | 0.5 +/- 0.3 | -51.3 +/- 2.7 | -324.9 +/- 23.6 | -7.1 +/- 3.6 | 0.7 +/- 0.6 | 1810.6 +/- 51.0 |
| 9 | -94.1 +/- 2.2 | 0.5 +/- 0.3 | -51.2 +/- 3.0 | -180.9 +/- 6.4 | -6.9 +/- 1.2 | 1.1 +/- 0.7 | 1496.6 +/- 44.7 |
| 10 | -102.8 +/- 2.1 | 0.5 +/- 0.3 | -49.9 +/- 1.2 | -250.5 +/- 23.4 | -2.9 +/- 2.3 | 0.9 +/- 0.6 | 1762.8 +/- 41.4 |
| 11 | -106.0 +/- 4.2 | 0.5 +/- 0.3 | -55.3 +/- 2.8 | -230.0 +/- 14.9 | -4.8 +/- 1.8 | 0.8 +/- 0.4 | 1744.7 +/- 54.6 |
| 12 | -112.3 +/- 1.6 | 0.5 +/- 0.3 | -60.0 +/- 1.9 | -267.3 +/- 16.4 | 1.1 +/- 1.0 | 0.4 +/- 0.1 | 1684.1 +/- 20.3 |
| 13 | -114.6 +/- 0.7 | 0.5 +/- 0.3 | -51.3 +/- 0.9 | -312.4 +/- 10.4 | -0.9 +/- 2.0 | 0.5 +/- 0.2 | 1643.4 +/- 36.6 |
| 14 | -100.0 +/- 1.2 | 0.5 +/- 0.3 | -58.4 +/- 1.0 | -152.4 +/- 11.9 | -11.2 +/- 2.0 | 0.6 +/- 0.2 | 1476.6 +/- 43.6 |
| 15 | -101.1 +/- 1.8 | 0.5 +/- 0.3 | -50.4 +/- 0.6 | -255.2 +/- 14.2 | 0.3 +/- 1.1 | 0.4 +/- 0.1 | 1681.2 +/- 26.8 |

**Table 5.** HADDOCK results for TLR4-SLP molecular docking (after refinement).

| Peptide # | HADDOCK score | RMSD | Van der Waals energy | Electrostatic energy | Desolvation energy | Restraints violation energy | Buried Surface Area |
| --- | --- | --- | --- | --- | --- | --- | --- |
| 1 | -123.1 +/- 1.7 | 0.5 +/- 0.3 | -64.9 +/- 3.3 | -166.3 +/- 1.8 | -27.7 +/- 0.6 | 1.6 +/- 0.3 | 1325.1 +/- 36.2 |
| 2 | -114.6 +/- 1.3 | 0.5 +/- 0.3 | -70.0 +/- 7.1 | -113.1 +/- 29.7 | -27.2 +/- 2.4 | 1.4 +/- 0.6 | 1390.1 +/- 27.5 |
| 3 | -134.0 +/- 7.8 | 0.5 +/- 0.3 | -67.2 +/- 2.1 | -208.0 +/- 13.0 | -22.4 +/- 3.1 | 1.0 +/- 0.2 | 1650.9 +/- 25.3 |
| 4 | -124.5 +/- 0.8 | 0.5 +/- 0.3 | -53.9 +/- 2.5 | -61.8 +/- 3.4 | -45.0 +/- 2.1 | 1.1 +/- 0.1 | 1427.0 +/- 24.6 |
| 5 | -99.8 +/- 2.0 | 0.5 +/- 0.3 | -49.3 +/- 1.6 | -101.3 +/- 15.5 | -25.8 +/- 1.0 | 1.0 +/- 0.1 | 1202.0 +/- 18.4 |
| 6 | -115.8 +/- 1.2 | 0.5 +/- 0.3 | -73.7 +/- 1.6 | -185.1 +/- 2.6 | -29.6 +/- 1.1 | 1.2 +/- 0.6 | 1365.7 +/- 47.1 |
| 7 | -108.9 +/- 0.4 | 0.5 +/- 0.3 | -52.6 +/- 1.7 | -60.7 +/- 13.5 | -23.2 +/- 1.6 | 1.0 +/- 0.2 | 1496.6 +/- 27.9 |
| 8 | -130.1 +/- 1.3 | 0.5 +/- 0.3 | -52.1 +/- 2.7 | -213.0 +/- 13.1 | -35.0 +/- 2.1 | 0.9 +/- 0.3 | 1334.6 +/- 21.8 |
| 9 | -93.1 +/- 1.9 | 0.5 +/- 0.3 | -52.6 +/- 1.5 | -141.9 +/- 20.9 | -12.7 +/- 2.8 | 1.8 +/- 0.9 | 1093.1 +/- 7.0 |
| 10 | -115.9 +/- 2.2 | 0.5 +/- 0.3 | -61.4 +/- 1.7 | -177.9 +/- 3.3 | -27.9 +/- 0.4 | 1.1 +/- 0.1 | 1336.5 +/- 30.6 |
| 11 | -114.6 +/- 0.9 | 0.5 +/- 0.3 | -37.2 +/- 1.9 | -79.0 +/- 3.6 | -37.5 +/- 2.9 | 0.8 +/- 0.2 | 1545.8 +/- 27.2 |
| 12 | -82.3 +/- 0.8 | 0.5 +/- 0.3 | -64.7 +/- 1.1 | -163.2 +/- 5.8 | -12.5 +/- 0.8 | 0.7 +/- 0.2 | 1080.6 +/- 25.5 |
| 13 | -115.2 +/- 1.2 | 0.5 +/- 0.3 | -62.2 +/- 1.3 | -96.5 +/- 7.5 | -31.2 +/- 0.6 | 0.8 +/- 0.1 | 1327.9 +/- 12.1 |
| 14 | -114.7 +/- 1.0 | 0.5 +/- 0.3 | -63.8 +/- 1.5 | -148.5 +/- 10.1 | -22.9 +/- 1.7 | 0.9 +/- 0.2 | 1380.8 +/- 28.2 |
| 15 | -114.0 +/- 1.7 | 0.5 +/- 0.3 | -62.2 +/- 1.1 | -153.0 +/- 8.0 | -19.8 +/- 0.4 | 2.2 +/- 0.7 | 1530.8 +/- 34.4 |

**Table 6.** Predicted ΔG and Kd for the TLR2/4-SLP interactions using PRODIGY.

| Peptide # | $\Delta G$ (kcal/mol) for TLR2 | Kd (M) for TLR2 | $\Delta G$ (kcal/mol) for TLR4 | Kd (M) for TLR4 |
| --- | --- | --- | --- | --- |
| 1 | -12.5 | $1.40 \times 10^{-9}$ | -9.5 | $2.10 \times 10^{-7}$ |
| 2 | -11.8 | $4.80 \times 10^{-9}$ | -9.9 | $1.10 \times 10^{-7}$ |
| 3 | -12 | $3.60 \times 10^{-9}$ | -11.5 | $8.00 \times 10^{-9}$ |
| 4 | -12.1 | $2.90 \times 10^{-9}$ | -10.2 | $6.70 \times 10^{-8}$ |
| 5 | -11.2 | $1.30 \times 10^{-8}$ | -9.3 | $2.70 \times 10^{-7}$ |
| 6 | -12.7 | $1.10 \times 10^{-9}$ | -9.5 | $2.10 \times 10^{-7}$ |
| 7 | -12.5 | $1.60 \times 10^{-9}$ | -10.3 | $5.60 \times 10^{-8}$ |
| 8 | -10.6 | $3.50 \times 10^{-8}$ | -10 | $8.70 \times 10^{-8}$ |
| 9 | -11.2 | $1.20 \times 10^{-8}$ | -10 | $8.90 \times 10^{-8}$ |
| 10 | -12 | $3.60 \times 10^{-9}$ | -11.2 | $1.30 \times 10^{-8}$ |
| 11 | -11.2 | $1.30 \times 10^{-8}$ | -11 | $1.80 \times 10^{-8}$ |
| 12 | -11.4 | $9.30 \times 10^{-9}$ | -9 | $4.90 \times 10^{-7}$ |
| 13 | -10.6 | $3.40 \times 10^{-8}$ | -12.3 | $2.10 \times 10^{-9}$ |
| 14 | -11.3 | $1.10 \times 10^{-8}$ | -8.6 | $8.20 \times 10^{-7}$ |
| 15 | -11.8 | $4.80 \times 10^{-9}$ | -10.4 | $4.30 \times 10^{-8}$ |

## Discussion

Despite the growing number of vaccine technologies, the COVID-19 pandemic is not over yet. By using the cancer-derived SLP technology, a new vaccine platform might be implemented against infectious diseases. SLPs function as a robust immune response trigger but they can also enhance the efficacy of the already available vaccines by providing an additional T cell epitope set.

Immunoinformatic approaches provide a rapid, cost-effective tool for designing antigenic molecules and explore their interactions with various effectors of the immune system.

This report shows how synthetic long peptides can be produced using in silico tools. Compared to other studies, the presented workflow uses a hybrid approach by exploiting data collected from COVID-19 convalescent patients.

A nineteen (15 class I and 4 class II) peptide pool was constructed using the data from a meta-analysis involving 852 COVID-19 patients worldwide. Not all populations are equally represented, leading to data being biased to certain favoured populations. To overcome this problem, further clinical studies are required to fully characterize the worldwide HLA haplotype occurrence and to construct a larger peptide database with epitopes identified from a larger cohort.

Designed synthetic long peptides comprised a class II-restricted epitope, a cathepsin-sensitive linker and a class I-restricted epitope. This design assures a bidirectional stimulation of both cellular and humoral immune responses with higher viral clearance.

SARS-CoV2 infection is associated with lymphocytopenia and an inversely proportional innate immune cells and cytokine increase with potential life-threatening effects. [35] Therefore, an immune priming with subsequent lymphocyte stimulation may be beneficial for preventing the evolution to severe COVID-19. [36] In addition, it was shown that active T cells mitigate the overly active innate immune response in mice, providing an additional benefit for T cell stimulation. [37]

Our vaccine design relies on intranasal administration via droplets. This formula is preferred because of its simplicity (can be administered by any individual, therefore reducing the number of medical professionals required for vaccination), tolerability (compared to traditional vaccination which some individuals find unpleasant or painful) and reduced number of complications. Additionally, intranasal vaccination provides mucosal immunity that targets viral particles right at the entry site. [39,40]

One of the major problems in peptide-based platforms involves potential allergenicity and toxicity. In silico studies have shown that the SLPs were neither toxic nor allergenic.

Protein folding using ab initio methods is an attractive method for solving tertiary structures due to their fairly rapid and cost-effective method compared to conventional protein analysis like cryo-EM or X-ray crystallography. ANN-based algorithms such as AlphaFold can replace wet lab protein structure solving in the future.

Toll-like receptors are pattern-recognition receptors (PRRs) involved in innate immune system antigen recognition. TLRs are horse-shoe-shaped transmembrane proteins which may be localized on the plasmalemma (recognizing extracellular pathogens) or inside the cytoplasm, attached to vesicles (recognizing intracellular microorganisms). TLR dimerization assures a higher recognition repertoire comprising fungal, bacterial or viral proteins. Recent studies demonstrated the involvement of TLR1, 2 and 6 in COVID-19-related cytokine storm by targeting the envelope protein. [36] Additionally, TLR4 was shown to be involved in an anti-bacterial-like early immune response by interacting with the SARS-CoV2 spike protein. [38] High-affinity interaction between TLR2/ 4 and the predicted synthetic long peptide pool suggests high immunogenicity and increased cytokine release, leading to subsequent internalization and antigen processing inside the dendritic cells. High similarities between the Gibbs free energies and dissociation constants result in an even distribution of the SLPs at the injection site and an equal binding probability to TLRs. Even though molecular docking showed positive results, molecular dynamic simulations are needed to check for the complex stability.

## Acknowledgments

The authors acknowledge the support of the OncoGen Association for making this research project possible.

